# New genetic codes in bacteria and archaea identified with a fast k-mer based algorithm

**DOI:** 10.64898/2026.04.02.715157

**Authors:** Artem V. Melnykov

## Abstract

The genetic code is conserved across all domains of life and is often described as universal. Nevertheless, many exceptions to the “universal” code have now been documented, most of these through manual or semiautomated inspection of highly conserved genes. Modern bioinformatics tools improved our ability to find alternative genetic codes but remain computationally expensive preventing widespread use on thousands of new species identified by sequencing environmental samples. Here I report a >100 fold accelerated method for inferring the genetic code directly from assembled genomes and apply it to thousands of previously uncharacterized assemblies from archaea and bacteria. I describe new candidate genetic code variations in both domains, including the first archaea sense codon reassignment. Identifying genetic code variations is important for understanding evolution of the standard code and improving accuracy of protein databases and open reading frame identification.

## Introduction

The set of rules for translating mRNA sequence into protein (the genetic code) is conserved in all domains of life. Mechanistically translational decoding is achieved by several interdependent RNA and protein factors all of which are individually highly conserved making any mutation in the genetic code extremely unlikely. In addition, the standard genetic code is believed to be optimized with respect to translation (1–3), so any deviation from the standard code is likely to be deleterious. Nevertheless, many variations have been documented over the years, starting with the observation of unusual code in human mitochondria (4). Since then, the majority of variant genetic codes have been found in organelles of eukaryotic cells but they also occur in nuclear genomes of unicelluar parasitic eukaryotes, fungi, and bacteria (reviewed in (5)). Documenting variations of the standard genetic code is important for understanding its evolution and for improving accuracy of protein databases and genome annotation tools.

Recent years have seen an explosion in the number of identified species of bacteria and archaea many of which belong to completely new phyla; the genetic codes used by them are unknown. Most of these new species were described based solely on their metagenome-assembled genomes (MAGs) and have never been grown in culture. Recovering their genetic codes requires a computational tool that can elucidate the code directly from the genome assembly.

The common approach to inferring the genetic code from a known genome is to rely on conservation of residues in proteins (6–10). First, the protein coding regions within the genome are identified followed by comparison of each predicted protein to the known consensus sequence. Mismatches in conserved amino acid residues serve as indicators of possible reassignments in the genetic code. Codetta, a systematic application of this approach with a sound theoretical foundation, was recently used to infer the genetic code from roughly 250,000 bacterial and archaeal genomes (10). While this tool is accurate and reliable, it is too slow to be routinely applied to thousands of newly sequenced prokaryotic genomes. The computational effort described in the original Codetta publication required a 30,000 core computing cluster.

Since then, the number of publicly available genomes increased at least tenfold putting database-wide application of Codetta out of reach for most researchers.

Here I present a new algorithm for inferring the genetic code from a genome assembly. This algorithm, like the methods mentioned above, is fundamentally based on conservation of amino acid residues in proteins. However, the search for protein coding regions at the heart of Codetta is replaced by lookup of short peptides (amino acid k-mers; other k-mer approaches in bioinformatics have been recently reviewed (11)) in a reference table representing conserved sequences in protein families. Once the conserved k-mers are identified, I apply the statistical model of Codetta to determine the likelihood of amino acid decoding for every codon. This alternative approach (from now on referred to as K-mer Assisted Code Inference, or KACI) leads to 144-fold speedup in the calculation required to infer the genetic code at the price of very slight decrease in accuracy compared to Codetta. I applied KACI to prokaryotic genome assemblies and confirmed all known nuclear codon reassignments as well as found new candidate reassignments in both bacteria and archaea.

## Results

### 1. Description of KACI

k-mer based representation of protein families is central to inferring the genetic code with KACI. This representation is accomplished by constructing a reference set of k-mers of fixed length for every protein family. In any given k-mer all positions except for one are occupied by amino acid residues and make up a short motif representative of the family. One position within this sequence is deemed uncertain, so a k-mer is accompanied by a set of probabilities describing the likelihood of finding a given amino acid at that position (Figure 1). In practice such k-mers and probabilities can be drawn from known protein sequences pre-grouped into families and processed as illustrated in Figure 1 for a hypothetical family A (see Methods for details). Briefly, all known proteins sequences from one family are cut up into overlapping k-mers of the same length. Those k-mers that differ at one amino acid position are then grouped together to form a reference k-mer with “?” at that position and a set of probabilities of finding a particular amino acid in this uncertain position. The process is then repeated for other protein families.

**Figure 1.**
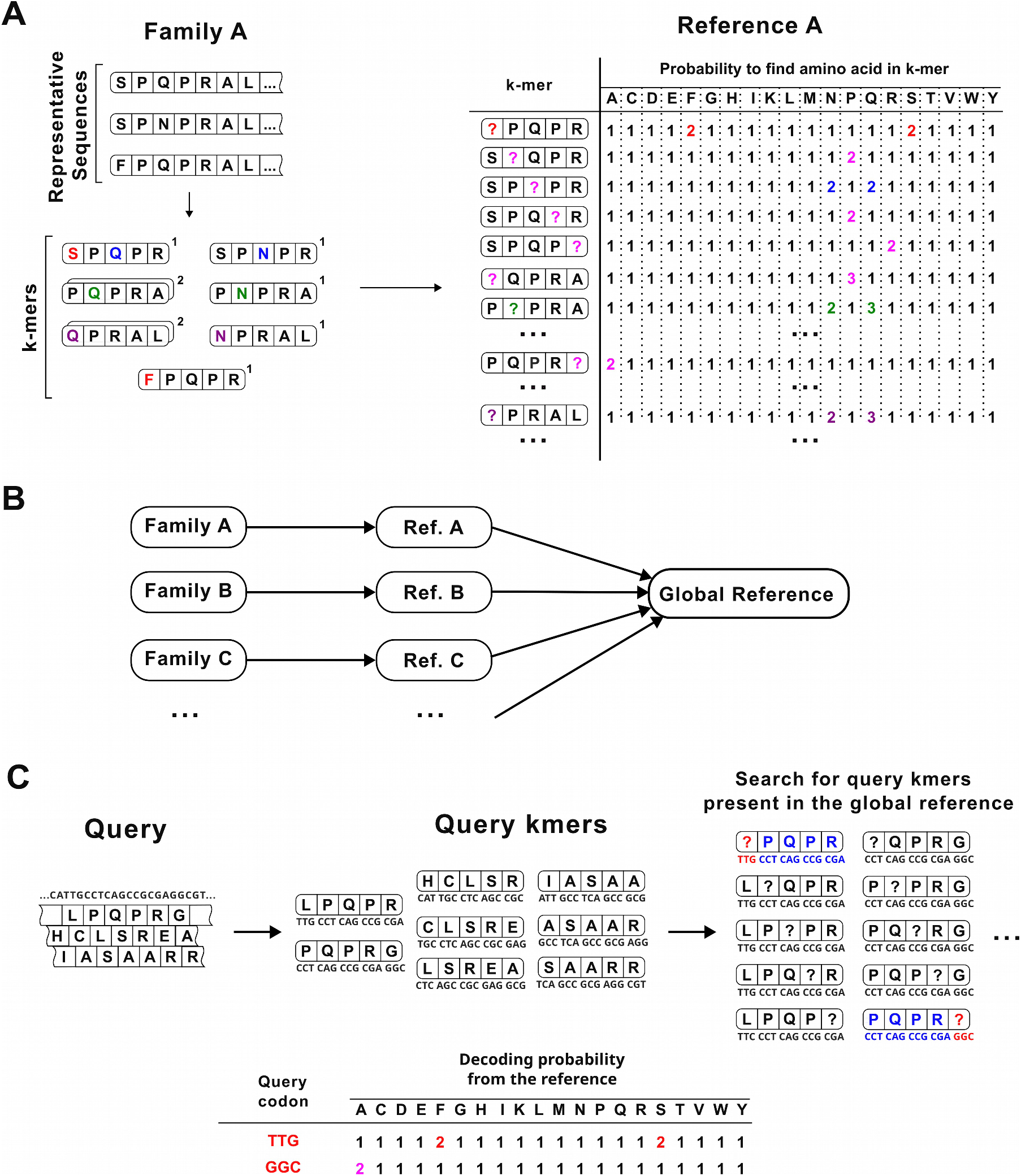
Illustration of the process used by KACI to construct the reference and infer decodings for a query. **A.** Reference construction for a hypothetical protein family based on the set of sequences available for that family. **B**. References from several protein families are combined into a single reference to be used for inferring the genetic code. **C**. Evaluation of a query by breaking up all possible translations into k-mers (only forward translations are shown for simplicity) and comparing these k-mers to the reference.

Finally, all k-mers are pooled together into a large reference where all protein families are represented. This reference can then be used to infer decoding probabilities of an unknown query (for example, a genome assembly).

Once the reference is constructed, the approach to inferring the genetic code from a genome sequence is very similar to the approach taken by Codetta. Each nucleotide sequence is translated into six open reading frames using the standard genetic code without stop codons (see Methods) as a reasonable guess. Every open reading frame is then broken up into overlapping k-mers of the same length as the reference k-mers. Then, for each position in a query k-mer I substitute the amino acid with the uncertainty symbol (“?”) and look for the matching k-mer in the reference. If a matching k-mer is found, the codon encoding the uncertain position is entered into the calculation of decoding probabilities based on the probabilities from the reference table. After pulling all probabilities together and appropriate normalization I obtain the decoding probability for every codon. The entire procedure mirrors that followed by Codetta with one important difference. In Codetta the probability of decoding a given codon instance comes from the HMM (hidden Markov model) profile aligned to that region. With KACI the probability comes from the k-mer that matches the sequence around the codon. Apart from the source of the decoding probabilities, KACI is identical to Codetta.

### 2. Evaluation of KACI

To evaluate the accuracy of KACI, I analyzed more than 200,000 NCBI genbank genomes for which Codetta results are available. Overall, the agreement between KACI and Codetta was excellent (Figure 2A and B) with 99.85% identical inferences for sense codons although KACI was slightly less sensitive (0.13% of Codetta inferences could not be confidently assigned by KACI). The two approaches gave different answers for 0.0045% of sense codons; the vast majority of these were attributed to the known artifact of Arg to Lys reassignment discussed in detail below. The agreement is slightly poorer for stop codons with 99% identical results. About 0.8% of the remaining stop codon inferences are cases where Codetta assigned a sense meaning to the codon but KACI did not. However, since neither approach is able to infer stop codons directly, the results for stop codons are prone to artifacts that I discuss in more detail below. Finally, it is worth noting that KACI successfully identified all known cases of genetic code reassignment in bacteria as detailed in the next section. Based on these findings, KACI provides accurate results with sensitivity only slightly lower than Codetta.

**Figure 2.**
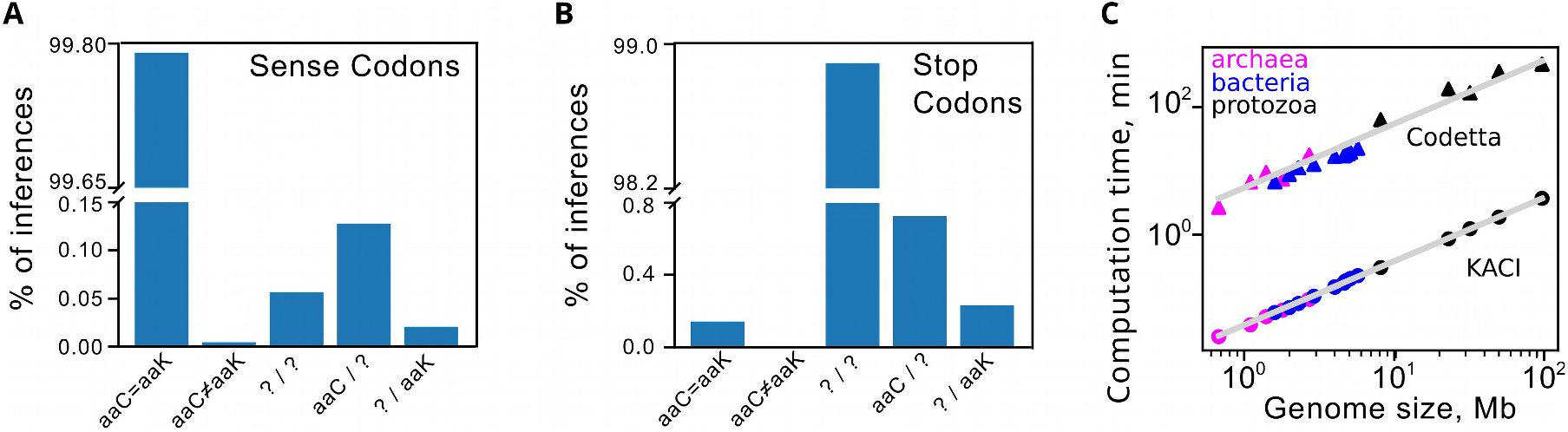
Comparison between KACI and Codetta. **A.** Percent of sense codons for which KACI and Codetta infer the same amino acid (aaC=aaK), different amino acids (aaC≠aaK), inference is uncertain for KACI (aaC / ?), Codetta (? / aaK) or both (? /?). **B**. Same as **A** but for stop codons (TAA, TAG, TGA) of the standard code. **C**. Computation time required to infer the genetic code for several representative genomes by Codetta and KACI.

The advantage of KACI is its reduced computational load. This concept is illustrated in Figure 2C where the computation time for inferring the genetic code is compared for genomes from bacteria, archaea, and eukaryota (protozoa only) that span genome sizes for organisms from these groups. While Codetta computation time scaled less predictably with genome size, in all cases KACI was between 100x and 200x faster (144x faster on average). This substantial improvement in analysis speed with very minor sacrifice in sensitivity allows processing very large volumes of data on a personal computer.

### 3. Analysis of bacterial and archaeal genomes

To search for new genetic codes, I analyzed approximately 2,700,000 bacterial and archaeal genome assemblies available at NCBI in July of 2025. All known or suspected codon reassignments (5) were successfully identified by KACI, namely, i) CGG reassignment from arginine to glutamine or tryptophan in several genera in the class Clostridia (10); ii) CGG and CGA reassignments from arginine to tryptophan in Absconditobacteria (10); iii) AGG reassignment from arginine to methionine in a clade within the genus Enterosoma (10); iv) TGA reassignment from a stop codon to glycine in the class JAEDAM01 (12, 13); v) TGA reassignment from a stop codon to tryptophan in the order Mycoplasmatales and several other clades of bacteria (14) (here and in the rest of the paper all classifications of prokaryotes are based on GTDB taxonomy (15)). In addition, I identified many new candidate genetic code reassignments three of which are supported by additional evidence and are discussed below (full inference results are available as a separate dataset (16)).

### 4. ACA reassignment from threonine to aspartate in bacteria

A new candidate codon reassignment of ACA from threonine to aspartate was identified in over 30 bacterial genome assemblies isolated across the world from soil and mine drainage samples. Approximately half of these assemblies are absent from the GTDB taxonomy because they failed quality check while those that are present belong to three genera of the family RAAP-2. The phylogenetic tree for a part of this family is shown in Figure 3; the close evolutionary relationship of all assemblies with this predicted reassignment provides additional evidence for its accuracy. Furthermore, most of the KACI predictions for these assemblies are confirmed by Codetta. Another piece of supporting evidence for this reassignment comes from the tRNA_UGU_ sequence which is rather different in the ACA reassignment clade and lacks the classic G1:C72 closing base pair associated with the Thr-tRNA identity (17, 18) (it is replaced by a weaker G:U base pair indicated by the red arrow in Figure 3B). Finally, the new meaning of ACA is illustrated with the alignment of a highly conserved sequence of COX1, subunit I of cytochrome c oxidase, where the conserved aspartate residue participates in binding a magnesium ion (19).

**Figure 3.**
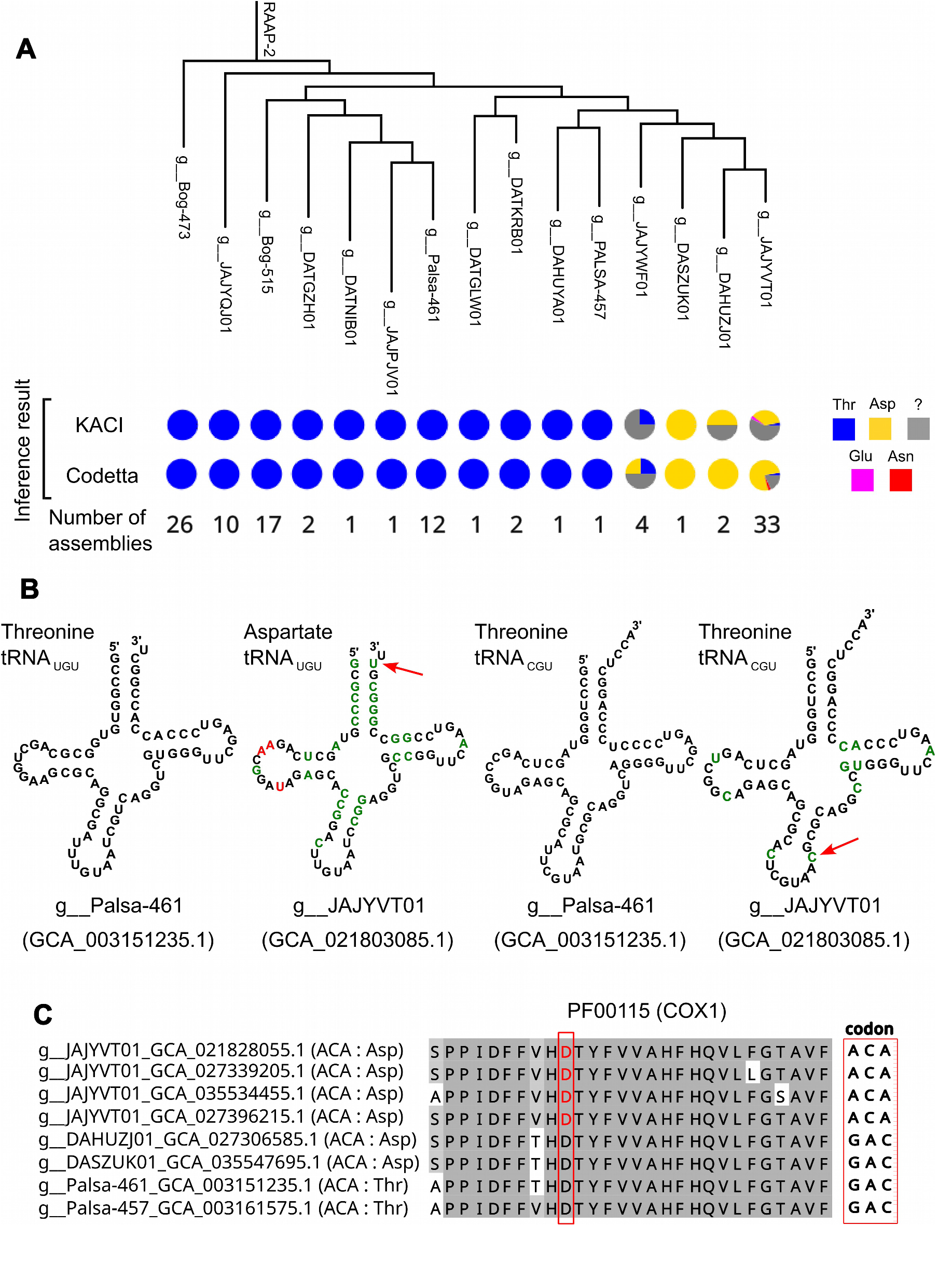
Reassignment of ACA codon from threonine to aspartate in the family RAAP-2. **A.** The phylogenetic tree of the portion of RAAP-2 including the ACA aspartate reassignment clade. Next to the tree are proportions of inferences for each genus as determined by KACI and Codetta. **B**. Representative secondary structures of tRNA_UGU_ and tRNA_CGU_ for genera JAJYVT01 (ACA reassignment clade) and Palsa-461 (outgroup). Mutations are show in green, insertions in red. The mutation of the threonine identity element (G1:C72 base pair) and the broken base pair in the acceptor stem are highlighted with red arrows. **C**. Alignment of a sequence motif from the catalytic core subunit of cytochrome C for several assemblies from the reassignment clade and the outgroup. The aspartates encoded by the reassigned ACA codon are shown in red.

All genome assemblies in the family RAAP-2 are characterized by a high GC content of 60-70%. It is very likely that this bias drove down the frequency of the ACA codon to the point where a change in its identity had little effect on protein translation, one of the hypothesized mechanisms of genetic code reassignment (20). The origin of the new Asp-tRNA_UGU_ is not clear as there is no obvious continuity of sequence or position relative to other genes between Thr-tRNA_UGU_ and Asp-tRNA_UGU_. It is possible that the original tRNA was lost and later replaced by horizontal gene transfer. Another interesting aspect of ACA reassignment is its potential effect on translation of ACG codons as tRNA_UGU_ could recognize ACG through a wobble base pair (21). It is unclear at this point what prevents such cross-recognition but it is worth mentioning that Thr-tRNA_CGU_ underwent a change in most of the assemblies predicted to decode ACA as aspartate: the closing base pair of the anticodon stem in these tRNAs is broken (Figure 3B). As this type of change in a tRNA is highly unusual it may be related to the reassigned meaning of the ACA codon.

### 5. CGG reassignment from arginine to alanine in bacteria

Another potential reassignment from arginine to alanine was found for CGG codon in 11 assemblies obtained from metagenome sequencing of human feces, human gut microbiome, and swine barn samples. Based on GTDB taxonomy, species with this reassignment form a clade in the genus RGIG3102 along with several bacteria species for which the inference for CGG was either uncertain or glutamine (Figure S4A). Many of the KACI results for this reassignments were not confirmed by Codetta which could not assign the decoding in several cases or assigned a different non-arginine meaning to this codon (KACI and Codetta do agree in assigning CGG to alanine in two assemblies not included in GTDB database). Although the new meaning of CGG is somewhat uncertain, CGG reassignment in this clade is supported by the tRNA_CCG_ sequences that lack the canonical A20 Arg identity element and possess the G3:U70 pair usually present in Ala-tRNAs (17, 18) (Figure S4B). This putative Ala-tRNA_CCG_ is also characterized by the closing G1:C72 base pair rather than G1:U72 found in the outgroup Arg-tRNA_CCG_ (the closing base pair of the tRNA often contributes to aminoacylation specificity). Notably, the only assembly in the suspected CGG reassignment clade that is predicted to decode CGG as arginine also has the conventional Arg-tRNA_CGG_, possibly because it was reacquired after its initial loss. Examples of protein sequence alignments (Figure S4C) lend further support to this reassignment. The genus RGIG3102 belongs to the class Clostridia in which several genera were predicted by Codetta to decode CGG as glutamine or tryptophan (10). Genomes assemblies in these genera are characterized by low GC content thought to favor reassignment by driving down the frequency of CGG codon among others. I tentatively add CGG reassignment to alanine to the list and note that many assemblies from Clotridia have uncertain CGG inferences accompanied by tRNA_CCG_ lacking arginine identity elements. Interestingly, tRNA_UCG_ is absent from these assemblies raising the question which tRNA is used for translating CGA (which is inferred to have its traditional arginine meaning). The extent of nonstandard CGG decoding and its exact meaning in Clostridia should be a subject of a separate investigation.

### 6. Genetic code changes in archaea

I also analyzed all archaea genome assemblies available at NCBI genbank. Most deviations from the standard genetic code identified in this dataset by KACI can be attributed to contamination by sequences of other species (such as CTG reassignment to serine, TAA/TAG reassignment to glutamate, and TGA to glycine or tryptophan all of which are well known in yeast and bacteria). The only known codon reassignments in archaea are context-dependent TAG coding for pyrrolysine and selenocysteine. As detailed in the Methods section, I did not include selenocysteine or pyrrolysine in the reference and therefore did not infer these decodings for any of the archaea assemblies. However, there are two assemblies (GCA_027068385.1 and GCA_964414255.1, both from samples isolated from marine hydrothermal vents) for which KACI inferred a sense codon change: CGG reassignment from arginine to tryptophan, a result that I confirmed by Codetta. One of these assemblies also contains a tRNA_CGG_ sequence that lacks an arginine identity element A20 and features an unusual bulge in the acceptor stem when compared to the conventional Arg-tRNA_CCU_ from the same assembly (Figure 4A). Further investigation into this reassignment is complicated by the fact that only one of these assemblies is currently present in the GTDB taxonomy, and it lacks any close relatives (although a preliminary assessment based on the percentage of amino acid identity for core genes suggests that GCA_027068385.1 and GCA_964414255.1 belong to the same family in the order Njordarchaeales, Figure S5). It is possible that this reassignment is an artifact of contamination by bacterial sequences for which this type of change of CGG meaning has been documented in Clostridia (10). However, as discussed below, sense codon inferences are less prone to contamination artifacts than stop codons. In addition, both KACI and Codetta identify CGG as the dominant tryptophan codon in these assemblies which is not typical for Clostridia. Finally, the inference for CGG is based partially on several moderately conserved tryptophan residues in proteins unique to archaea and eukaryotes, such as ribosomal proteins L32e and S27 (Figure 4B). Based on these observations I conclude that the inferred CGG reassignment to tryptophan is accurate and is specific to the archaea assemblies.

**Figure 4.**
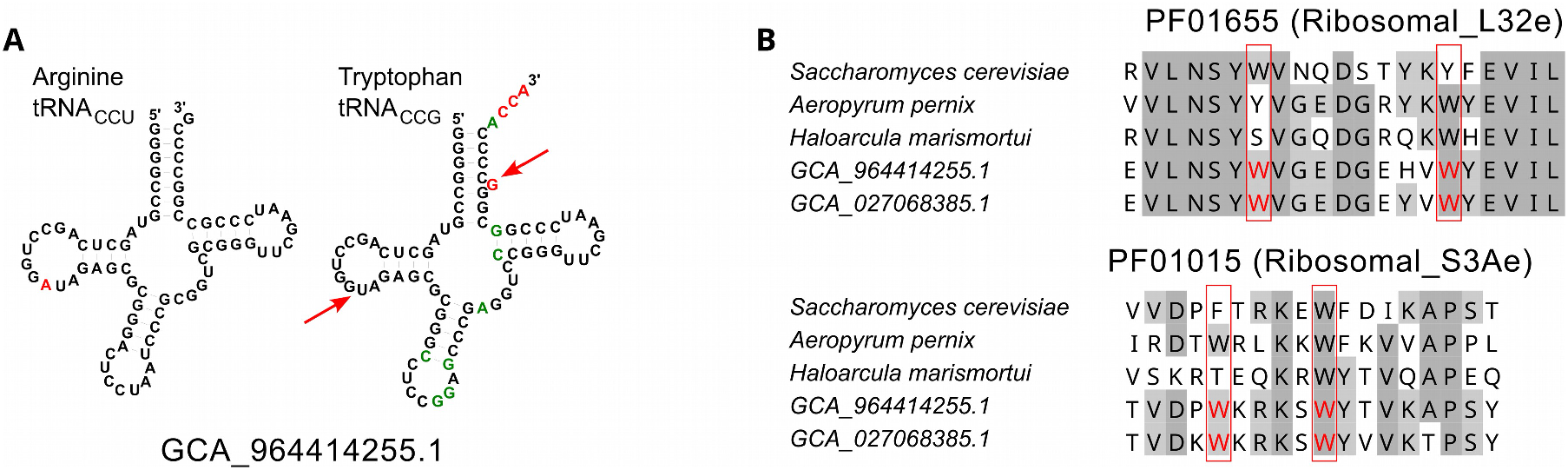
Reassignment of CGG codon from arginine to tryptophan in two archaea genome assemblies. **A.** Secondary structures of tRNA_CCU_ and tRNA_CCG_ from GCA_964414255.1. The absent arginine identity element A20 and the unusual acceptor stem bulge in tRNA_CCG_ are shown with red arrows. **B**. Alignment of short sequence motifs from two ribosomal proteins characteristic of eukaryota and archaea but not bacteria. The motifs are compared to sequences from budding yeast and two model archaea. The tryptophans encoded by CGG are shown in red.

The reassignment of CGG in archaea is different from the known change in this codon in bacteria. First, both archaea assemblies are characterized by balanced GC content (around 50%), so this codon reassignment was either not driven by the low frequency of the CGG codon or the GC content rebalanced since the initial change in the codon meaning. Second, unlike Clostridia, both GCA_027068385.1 and GCA_964414255.1 assemblies are characterized by complete absence of CGA codons in sequences that can be identified as known HMM domains. As a result, Codetta (and KACI) cannot assign a meaning to CGA raising the possibility that it is in fact a stop codon. Interestingly, the cognate tRNA_UCG_ is not found in either of the assemblies. Unfortunately, the approach taken by Codetta and KACI does not allow direct inference of stop codons. The question of CGA meaning in these archaea will require a separate investigation.

### 7. Limitations

The analysis of prokaryotic genome assemblies also identified several cases in which KACI can produce inaccurate results. First, there is a small fraction of bacterial genomes in which the inferred meaning of one of the arginine codons (usually AGG) is lysine. Unlike other candidate codon reassignments, this result is not supported by any additional evidence and I believe that these inferences are an artifact of the approach. Codetta also suffers from the same artifact (10); it is likely that insufficient representation of some proteins in the reference database leads to overestimation of lysine decoding probability in positions where arginine is well tolerated. For now, any cases of predicted arginine codon reassignment to lysine should be interpreted with extra caution.

In several cases KACI produced an inference for a sense codon that was not confirmed by Codetta and was not substantiated by other lines of evidence. I found that in most of these cases the proportion of “end” k-mers (k-mers with “?” at one of the ends) was unexpectedly high, sometimes approaching 100%. Since there is no reason why end k-mers should be overrepresented among the k-mers used to produce the inference, any inference with a high fraction of end k-mers may not reliable. I found that many such k-mers do not map to predicted open reading frames and likely reflect presence of noncoding (but possibly open reading frame-derived) sequences in bacterial genomes.

Another potential source of error in genetic code inference is contamination of assembled MAGs with sequences from other species that do utilize unusual genetic codes. This is especially true for well established codon reassignments such as CTG reassignment from leucine to serine in yeast and of stop codons TAA and TAG to glutamine in protista. Any genomes contaminated with such sequences may produce erroneous results, especially if the codon is naturally characterized by low abundance in coding sequences (which is always true for TAA and TAG in species that use the standard genetic code). Any predicted inference, especially sense meaning of the traditional stop codons, should be supported by additional evidence such as species phylogeny that is consistent with the reassignment and presence of the cognate tRNAs with unique characteristics. Ultimately, only experimental evidence can firmly establish a variation in the genetic code.

## Discussion

In this report I described an accelerated algorithm for inferring the genetic code used by an organism based on sequences from its genome. While similar in many ways to another tool (Codetta), KACI is considerably faster because it looks up short sequence motifs from a reference table instead of relying on the time consuming sequence alignment step. This gain in computation speed is offset by somewhat lower accuracy. However, as I showed here, a typical genome assembly contains more than enough information for KACI to accurately infer the genetic code. One of the reasons for KACI’s good performance may be its ability to capture high order amino acid position correlations within a short sequence motif.

After applying KACI to prokaryotic genome assemblies I found three new sense codon reassignments: ACA threonine to aspartate, CGG arginine to alanine (both in bacteria), and CGG arginine to tryptophan in archaea. Reasignment of arginine codons so far is the most common kind of nuclear genetic code variation in prokaryotes. On the other hand, the change in the threonine codon identity is, to my knowledge, the first ever for this amino acid in nuclear or organellar genomes. The archaea CGG reassignment identified here is also interesting not only because it is the first example of a sense codon change in this domain of life but also because it may be accompanied by a sense (CGA) to stop codon change. Future investigation and ideally experimental verification will shed more light on the genetic code variations reported here. In the meantime, KACI should become a valuable tool for analyzing vast amounts of sequencing data for newly identified species and suggesting which might adopt nonstandard genetic codes. It is almost certain that more genetic code variants remain to be discovered.

## Materials and Methods

### Reference construction, query evaluation, and the choice of reference parameters

A custom library implementing KACI was used to construct the reference and evaluate queries (it is available at https://github.com/artmeln/lib-kaci). The reference was constructed from the compiled 90% non-redundant set of all protein sequences (Pfam-A.fasta.gz) that accompanied the release of v.37.2 of the Pfam database (22) (in processing these protein sequences, only the 20 universal amino acids were allowed, and selenocysteine (U) and pyrrolysine (O) were treated as breaks in the sequence). As illustrated in Figure 1, the protein sequences from one family were broken up into overlapping k-mers of specified length. At this point any k-mer that contained 4 or more instances of the same amino acid was discarded (this step reduces the contribution of amino acid repeats); k-mers that were encountered only once in the family were also discarded. In the next step, for each position in the k-mer the probability estimate for finding a specific amino acid at that position was calculated by adding the number of times each amino acid was observed to the baseline count of 1. The k-mer with one uncertain position which was obtained through this procedure was only kept if the number of k-mers contributing to the probability estimate was above a certain threshold which I call link number (this step limits the reference to the sequence motifs with a higher degree of conservation and makes the size of the final reference more manageable). Once the reference for one family is constructed through this process, it is added to the global reference. If at this point it is discovered that a certain k-mer is present in more than one family, it is masked and not used in query estimates. Finally, the procedure is repeated for all protein families. Once the reference is constructed, a query (a genome assembly sequence) was processed according to the procedure described in the main text (Figure 1C). To limit the contribution of short sequence repeats, the number of times a query k-mer was allowed to contribute to the calculation of the decoding probability was limited to 2. Any decoding that was based on fewer than 4 k-mers or had the decoding probability below 0.9999 was considered uncertain.

The two main parameters in reference construction are the length of the k-mer and the link number. To choose the optimal k-mer length, I analyzed 1,000 genomes selected at random from the dereplicated set of bacterial genomes analyzed with Codetta and predicted to use the standard genetic code. The inference was carried out for several k-mer lengths and link numbers; the results shown in Figure S1 suggest that k=11 is optimal, and this k-mer length was kept for all further calculations. Next, I evaluated the effect of using a translation table without stop codons in the preliminary translation of the genome sequence on the inference of sense codons (using a table without stop codons in this step is important for accurately inferring the meaning of these codons in genomes that do use them for amino acid coding). As shown in Figure S2, using the translation table with TAA-TAG-TGA coding for Ala-Gly-Ser or Gly-Leu-Thr has minimal disturbing effect on the inferred meaning of sense codons. The Ala-Gly-Ser combination was used in the rest of the calculations.

Finally, the same dataset of 1,000 genomes was used to infer codon meanings of both sense and stop codons using the just discussed translation table without stop codons as a function of link number. As shown in Figure S3, the choice of the link number is a trade-off between increasing the number of correctly inferred sense codons and reducing the number of stop codons with incorrectly assigned sense meaning. For the rest of the calculations I chose link number 20 to balance these factors. Another reason for choosing a larger link number is to keep the size of the reference smaller. With k=11 and link number=20 chosen here, the size of the reference when stored on hard drive is 4 GB, and the amount of RAM required for loading this reference is approximately 15 GB.

### Comparison to Codetta

All calculations were carried out on a workstation with a Xeon E5-2699 v3 processor. In order to run KACI, it is necessary to load a precalculated reference table into memory, a step that for the reference parameters listed above takes a little less than a minute. This step was not included while timing KACI performance since in a typical application the reference table is loaded once followed by many KACI inferences for genome assemblies run in parallel as well as sequentially.

### Bioinformatics tools

tRNAs in genome assemblies were identified with tRNAscan-SE (v.2.0.12) (23) and aligned with Clustal W (v.2.1) (24). The secondary structures of the tRNAs were visualized using R2DT (25). The phylogenetic trees were constructed from GTDB taxonomy database (version RS226)(15) and visualized using ETE (26). For archaea assemblies missing from the GTDB taxonomy, the amino acid identity was calculated using EzAAI (27) on the set of archaea core single copy genes (28) extracted from genome assemblies using anvi’o (29, 30).

## Notes

### Competing Interest Statement

The authors have declared no competing interest.

