## Supplemental Figures for "New genetic codes in bacteria and archaea identified with a fast k-mer based algorithm"

Artem V. Melnykov

### Figures

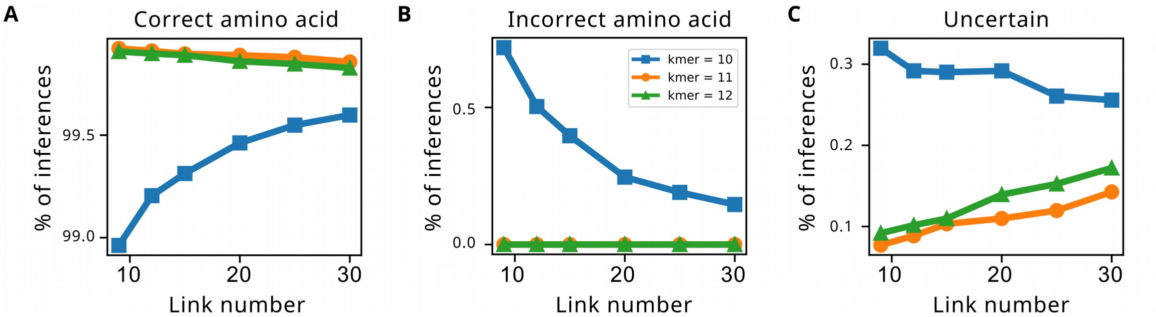


**Fig. S1.** Sense codon meanings inferred by KACI for a test set of 1,000 genomes **A.** correctly inferred amino acid; **B.** incorrectly inferred amino acid; **C.** uncertain inference. The results shown are for three kmer lengths (10, 11, 12 amino acids) and different link numbers (see Methods). The standard code was used for initial translation of the query.

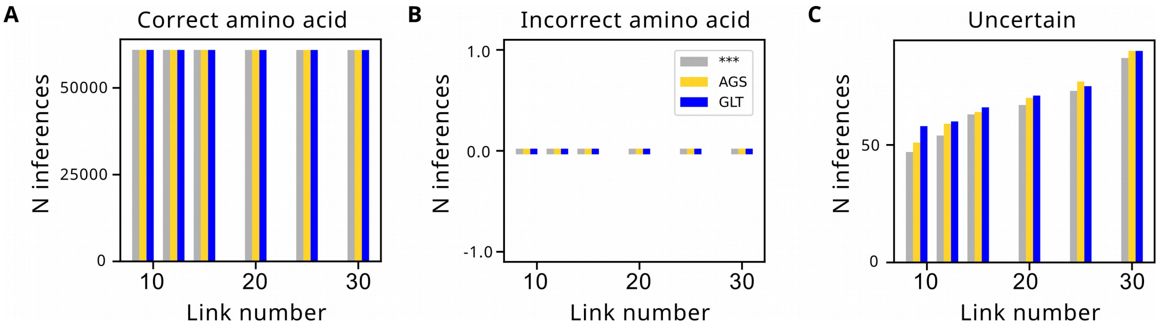


**Fig. S2.** Sense codon meanings inferred by KACI for a test set of 1,000 genomes using the standard code translation table (TAA=*, TAG=*, TGA=*: purple), and the standard table without stop codons (TAA=Ala, TAG=Gly, TGA=Ser: fuchsia; TAA=Gly, TAG=Leu, TGA=Thr: blue). **A.** correctly inferred amino acid; **B.** incorrectly inferred amino acid; **C.** uncertain inference. The results shown are for kmer length 11 and different link numbers (see Methods).

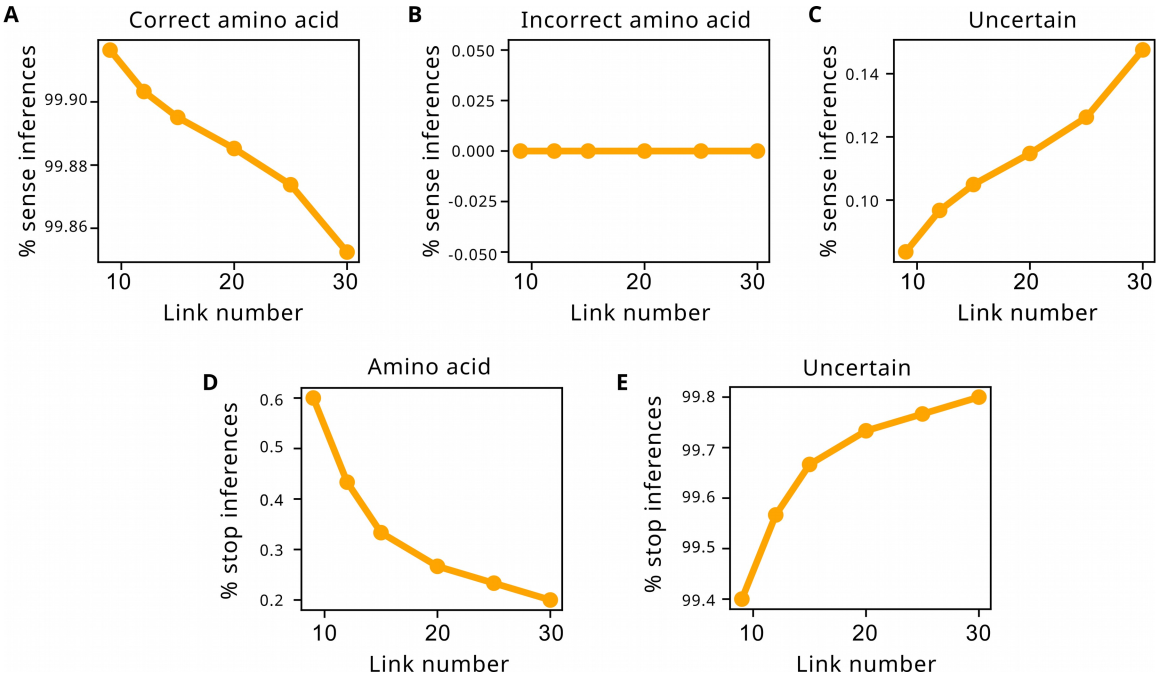


**Fig. S3.** Sense and stop codon meanings inferred by KACI for a test set of 1,000 genomes using the standard code translation table without stop codons (TAA=Ala, TAG=Gly, TGA=Ser). **A.** correctly inferred sense codon; **B.** incorrectly inferred sense codon; **C.** uncertain meaning inferred for sense codon; **D.** amino acid meaning inferred for stop codon; **E.** uncertain meaning inferred for stop codon.

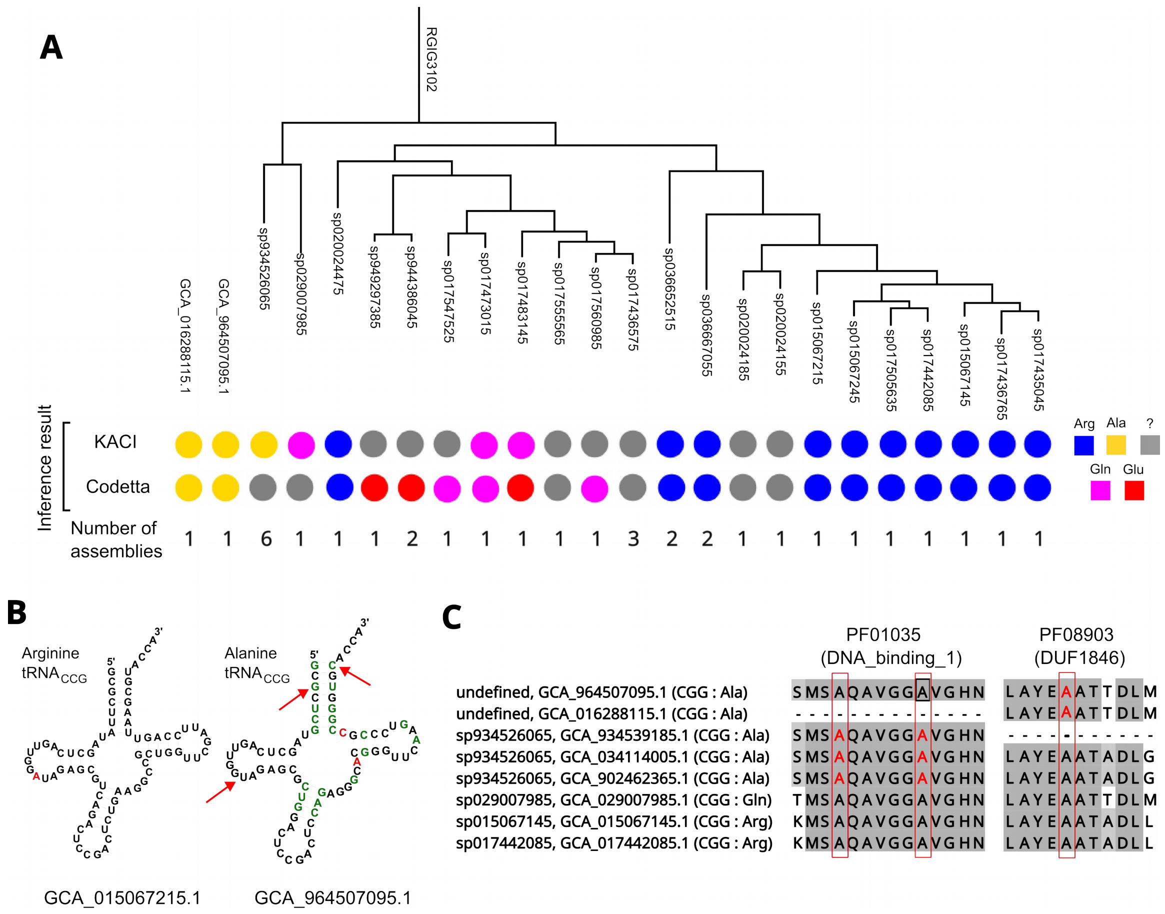


**Fig. S4.** Reassignment of CGG codon from arginine to alanine in the genus RGIG3102. **A.** The phylogenetic tree of RGIG3102. Next to the tree are proportions of inferences for each species as determined by KACI and Codetta. **B.** Representative secondary structures of tRNA-CCG for the CGG reassignment clade and for the outgroup. Mutations are show in green, insertions in red. The missing arginine identity element (A20), the mutated closing base pair (G1:U72) and the alanine identity element (G3:U70) are highlighted with red arrows. **C.** Alignment of two protein sequence motifs for several assemblies from the reassignment clade and the outgroup. The alanines encoded by the reassigned CGG codon are shown in red.

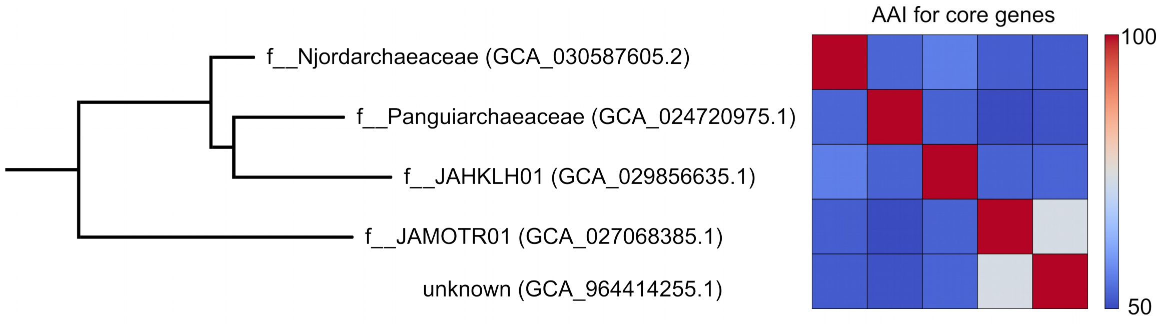


**Fig. S5.** The phylogenetic tree of the order Njordarchaeales. Next to the tree is the amino acid identity (AAI) matrix for the core single copy genes from each assembly. The relatively high AAI (75%) between GCA_027068385.1 and GCA_964414255.1 indicates that they might belong to the same family.
